# *Vibrio cholerae* OmpR represses the ToxR regulon in response to membrane intercalating agents that are prevalent in the human gastrointestinal tract

**DOI:** 10.1101/752626

**Authors:** DE Kunkle, TF Bina, XR Bina, JE Bina

## Abstract

Multidrug efflux systems belonging Resistance-Nodulation-Division (RND) superfamily are ubiquitous in Gram-negative bacteria. RND efflux systems are often associated with multiple antimicrobial resistance but also contribute to the expression of diverse bacterial phenotypes including virulence, as documented in the intestinal pathogen *Vibrio cholerae*, the causative agent of the severe diarrheal disease cholera. Transcriptomic studies with RND efflux-negative *V. cholerae* suggested that RND-mediated efflux was required for homeostasis, as loss of RND efflux resulted in the activation of transcriptional regulators, including multiple environmental sensing systems. In this report we investigated six RND efflux responsive regulatory genes for contributions to *V. cholerae* virulence factor production. Our data showed that *V. cholerae* gene VC2714, encoding a homologue of *Escherichia coli* OmpR, was a virulence repressor. The expression of *ompR* was elevated in an RND-null mutant and *ompR* deletion partially restored virulence factor production in the RND-negative background. Virulence inhibitory activity in the RND-negative background resulted from OmpR repression of the key ToxR regulon virulence activator *aphB*, and *ompR* overexpression in WT cells also repressed virulence through *aphB.* We further show that *ompR* expression was not altered by changes in osmolarity, but instead was induced by membrane intercalating agents that are prevalent in the host gastrointestinal tract, and which are substrates of the *V. cholerae* RND efflux systems. Our collective results indicate that *V. cholerae ompR* is an *aphB* repressor and regulates the expression of the ToxR virulence regulon in response to novel environmental cues.

## INTRODUCTION

The Gram-negative bacterium *Vibrio cholerae* is the causative agent of the life-threatening diarrheal disease cholera. *V. cholerae* is an aquatic organism and infects humans following the consumption of *V. cholerae* contaminated food or water. After ingestion *V. cholerae* colonizes the small intestine epithelium to cause disease by a process that is dependent upon virulence factor production. The two most important *V. cholerae* virulence factors are the toxin coregulated pilus (TCP), which mediates intestinal colonization, and cholera toxin (CT), an enterotoxin that is responsible for the secretory diarrhea that is the hallmark of the disease cholera (1). CT and TCP production are under the control of a hierarchical regulatory system known as the ToxR regulon (2). Activation of the ToxR regulon begins with expression of two cytoplasmic transcriptional regulators, *aphA* and *aphB* (3, 4). AphA and AphB function synergistically to activate *tcpP* expression. TcpP then binds along with ToxR to the *toxT* promoter to activate *toxT* expression. ToxT directly activates the expression of the genes that encode for CT and TCP production (2).

The expression of adaptive responses is important for the success of *V. cholerae* as a pathogen. This includes tight regulation of the ToxR regulon which is known to limit virulence factor production to specific niche within the host. Thus, the ToxR regulon has evolved to respond to specific environmental signals within the host (5). Other genes, which are important for survival and persistence in aquatic ecosystems, must be repressed during host entry for successful colonization (6-8). Late in infection, in preparation for host exit, *V. cholerae* downregulates virulence genes while activating genes required for dissemination and transmission (9-12). Although the genetic mechanisms involved in ToxR regulon activation have been extensively studied, little is known about how environmental signals influencing ToxR regulon expression *in vivo*.

*V. cholerae* is exposed to disparate environments in the aquatic ecosystem and the human gastrointestinal tract. *V. cholerae* survival and growth in these niches requires rapid adaptation to environmental conditions. *V. cholerae* enters humans from aquatic ecosystems that are typically aerobic and alkaline. The bacterium must then pass through the gastric acid barrier of the stomach before entering the duodenum and migrating to the epithelial surface where it colonizes the crypts of the small intestine. Successful transition between these dissimilar environments requires that *V. cholerae* modulate its transcriptional responses so that specific genes are only expressed in appropriate niches. In *V. cholerae*, like most bacteria, this is achieved by environmentally responsive regulatory systems that monitor the extracellular environment using a range of membrane bound sensors such as ToxR and two-component signal transduction systems (TCS) (13).

TCS are widespread phospho-relay systems that modulate gene expression in response to environmental cues. They consist of a membrane-bound histidine kinase sensor protein coupled with a cytosolic response regulator. In the presence of appropriate stimuli, the sensor auto phosphorylates a conserved histidine residue before transferring the phosphate to a conserved aspartate residue on the response regulator to activate the response regulator. Activated response regulators function to modulate adaptive responses by effecting the expression of target genes. Response regulators are typically transcription factors, but also can function by other mechanisms (14). The adaptive responses mediated by TCS are broad and include virulence, motility, metabolism and stress responses.

One of the better characterized TCS is the EnvZ-OmpR system that is ubiquitous in Gram-negative bacteria (15). EnvZ is the membrane associated sensor kinase and OmpR the response regulator that functions as a transcription factor. EnvZ-OmpR was first discovered in *Escherichia coli* and shown to regulate the expression of its two major outer membrane porin proteins (OMP), *ompC* and *ompF*, in response to environmental osmolarity (16-18). The function of OmpR as an osmoregulator has been extended to a number of other bacteria genera (19-21). OmpR has also been linked to other phenotypes in Gram-negative bacteria including virulence (19, 20, 22-26) and acidic tolerance (21, 27-31). The *V. cholerae* OmpR homologue (open reading frame (ORF) VC2714) has been poorly studied, and its role in *V. cholerae* biology is unknown.

The RND efflux systems are ubiquitous tripartite transporters in Gram-negative bacteria that play critical roles in antimicrobial resistance. Many RND efflux systems exhibit broad substrate specificity and have the capacity to efflux multiple substrates that are both structurally and functionally unrelated (32, 33). The RND systems play critical roles in antimicrobial resistance by exporting toxic compounds from the cytosol and periplasm into the extracellular environment. Although RND efflux pumps have been widely studied for their role in multiple antibiotic resistance, they also impact many other physiological phenotypes in bacteria (34). This was recently documented in *V. cholerae* where the RND systems were shown to be required for cell homeostasis (33, 35, 36). The absence of RND efflux in *V. cholerae* resulted in downregulation of the ToxR regulon and altered expression of genes involved in metabolic and environmental adaptation (37, 38), including several TCS. The results of these studies suggested that RND-mediated efflux modulated homeostasis by effluxing cell metabolites which served as concentration-dependent environmental cues to initiate transcriptional responses via periplasmic sensing systems. This observation suggested the possibility that these TCS may have contributed to the virulence attenuation observed in efflux impaired *V. cholerae*.

In this work we investigated six regulatory genes that were induced in the absence of RND-mediated efflux for their contribution to virulence factor production in *V. cholerae*. This revealed that VC2714, encoding a homolog of *E. coli* OmpR, functioned as a virulence repressor in *V. cholerae*. We documented that VC2714 repressed the expression of the key virulence regulator *aphB.* We further showed that *ompR* expression was regulated in response to detergent-like compounds which are prevalent in the host gastrointestinal tract and are substrates of the RND transporters. Our collective results suggest that the *V. cholerae* EnvZ-OmpR TCS has evolved to regulate virulence in response to novel environmental stimuli.

## RESULTS

### *V. cholerae ompR* represses virulence factor production

The loss of RND-mediated efflux resulted in downregulation of the ToxR regulon and diminished CT and TCP production (37), suggesting that there is one or more factors linking efflux to virulence factor production. Transcriptional profiling of an RND negative *V. cholerae* mutant during growth under AKI conditions (i.e. virulence inducing conditions) showed that the expression of a number of regulatory genes, including several TCSs, were increased in the absence of RND efflux (38). We hypothesized that one or more of these regulatory genes may have contributed to RND efflux-dependent virulence repression. To test this, we expressed six regulators (i.e. VC0486, VC1320*-*VC1319, VC1081, VC1638, VC1825, VC1320 and VC2714) from the arabinose-regulated promoter in pBAD33 in WT *V. cholerae* strain JB58 during growth under AKI conditions in the presence of 0.05% arabinose and quantified CT production. VC0486 encodes an uncharacterized DeoR family regulator. VC1320 (*carS*) and VC1319 (*carR*) encode the CarRS TCS that is involved in regulating LPS remodeling and *vps* production (39-41); *carR* (pTB15)and the *carRS* (pTB3) were independently expressed in *V. cholerae*. VC1081 encodes an uncharacterized response regulator. VC1638 was recently shown to regulate the expression of *vca0732* in response to polymyxin B (42). VC1825 is an AraC-family regulator that regulates a PTS transporter (43). VC2714 encodes an uncharacterized response regulator. The results showed that only pTB11, expressing VC2714, repressed CT production (Fig. 1A). VC2714 encodes a homolog of the *E. coli* osmotic stress regulator OmpR, with 92.1% amino acid sequence similarity, and hereafter will be referred to as *ompR*.

**Figure 1.**
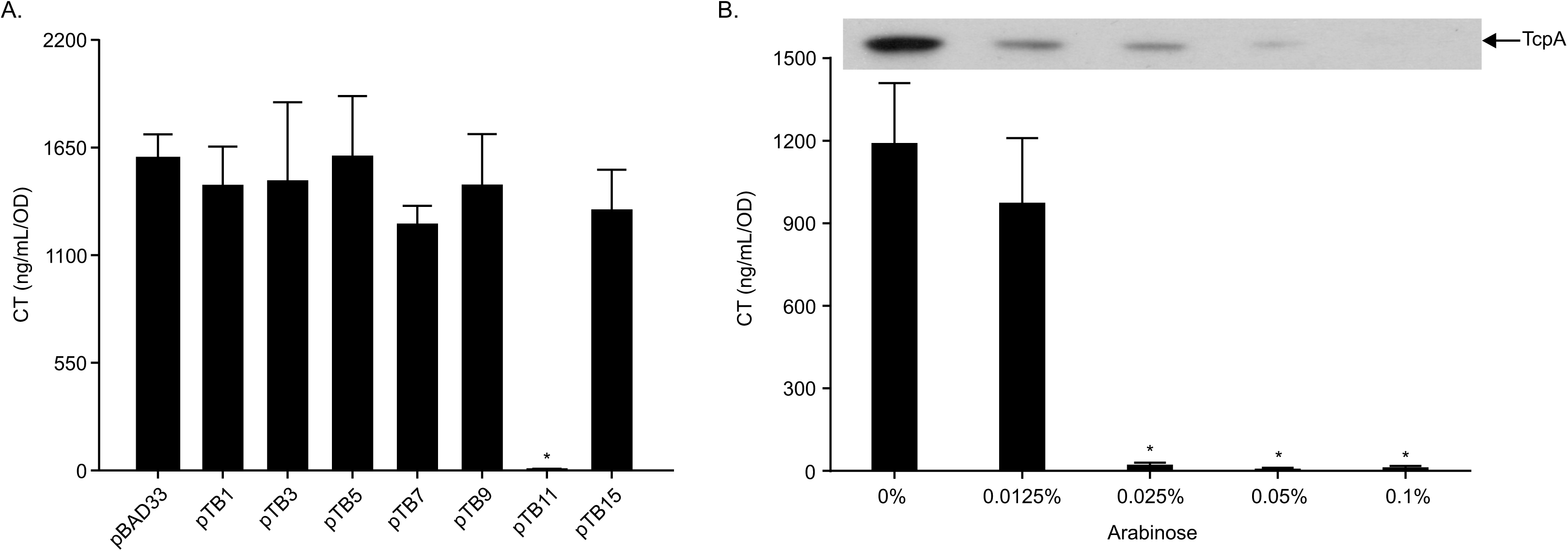
Overexpression of *ompR* represses virulence factor production. (A) WT *V. cholerae* harboring pBAD33 or the indicated expression plasmids were cultured under AKI conditions with 0.05% arabinose for 24h when culture supernatants were used for CT quantitation. Data represents the mean +/- SD of three independent experiments. * P<0.001 relative to pBAD33. (B) WT *V. cholerae* harboring pTB11 (pBAD33-*ompR*) was cultured under AKI conditions with indicated arabinose concentrations for 24h when culture supernatants were used for CT quantitation by GM1 ELISA and the cell pellets for TcpA immunoblotting, respectively. CT data represents the mean +/- SD of a minimum of three independent experiments. * P<0.001 relative to 0%. The TcpA immunoblot is representative of a minimum of three independent experiments.

To further verify that *V. cholerae ompR* was a virulence repressor we repeated the above experiment in WT strain JB58 harboring plasmid pTB11 during growth under AKI conditions in the presence of increasing arabinose concentrations and quantified CT and TcpA production. The results showed an arabinose-dependent inhibition of both CT and TcpA production (Fig. 1B). Based on these results we concluded that *ompR* functions as a virulence repressor in *V. cholerae*.

### OmpR contributes to virulence repression in RND-efflux deficient *V. cholerae*

To verify that *ompR* was upregulated in RND-deficient *V. cholerae* as previously indicated in a transcriptomics dataset, (38), we introduced the *ompR-lacZ* transcriptional reporter plasmid pKD9 into JB58 and the isogenic RND efflux-negative strain JB485, and quantified *ompR* expression in both strains following growth in LB broth, minimal T-media, and under AKI conditions. The results showed significantly increased *ompR* expression in JB485 relative to WT during growth under AKI conditions, but no significant difference in LB broth or minimal T-medium (Fig. 2A). These findings confirmed the previous study and suggested that the RND-efflux dependent induction of *ompR* transcription was specific to AKI growth conditions.

**Figure 2.**
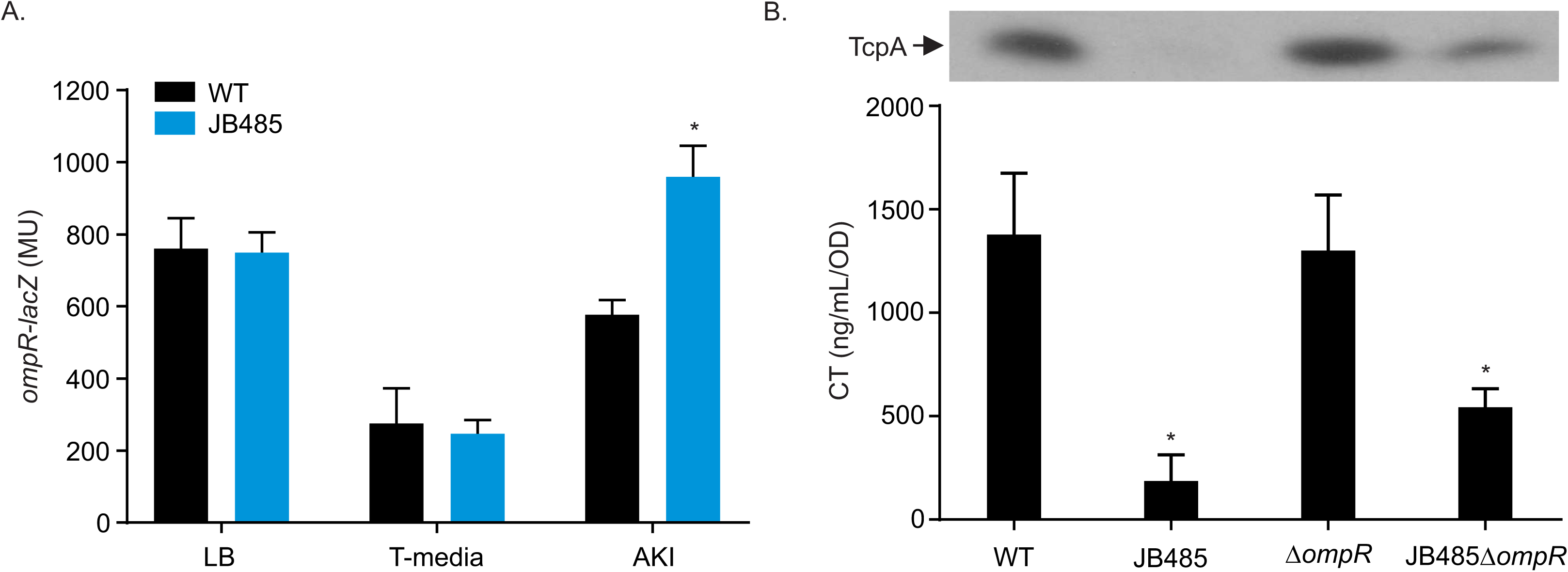
OmpR represses virulence factor production in RND efflux negative *V. cholerae*. (A) WT and JB485 (ΔRND) *V. cholerae* strains harboring an *ompR-lacZ* reporter plasmid were cultured under the indicated conditions for 5h when β-galactosidase activity was quantified. Data represents the mean +/- SD of three independent experiments performed in triplicate. * P<0.001 relative to WT. (B) The *V. cholerae* strains JB58 (WT), JB485 (ΔRND), DK243 (JB58Δ*ompR*) and DK246 (ΔRNDΔ*ompR*) were cultured under AKI conditions for 24h when CT and TcpA production were assessed by GM1 ELISA and TcpA immunoblotting, respectively. The CT data represents the mean +/- SD of a minimum of three independent experiments. * P<0.01 relative to the parental strain. TcpA immunoblot is representative of a minimum of three independent experiments.

We next tested if *ompR* contributed to the virulence repression observed in the RND-negative strain JB485. To address this, we created *ompR* deletion strains in WT JB58 and RND-negative strain JB485 and quantified CT and TcpA production in WT, JB485 and their respective isogenic Δ*ompR* mutants. Consistent with previous studies (37), the RND-negative strain produced significantly reduced amounts of CT and TcpA relative to WT (Fig. 2B) and deletion of *ompR* in WT did not significantly affect CT or TcpA production. By contrast, deletion of *ompR* in JB485 partially restored CT and TcpA production relative to wild type, but the magnitude of the increase did not reach WT levels (Fig. 2B). Together these data suggested that *ompR* contributed to virulence attenuation in the RND negative background, but that other factors are also involved in virulence repression.

### *V. cholerae* OmpR represses *aphB* expression

The above results suggested that OmpR was a virulence repressor, but the mechanism by which it attenuated virulence factor production was unclear. As CT and TCP production are positively regulated by the ToxR regulon, we hypothesized that OmpR repressed components of the ToxR regulon. If this was true, then *ompR* deletion in JB485 should increase the expression of the affected ToxR regulon genes, relative to the parental strain JB485. We therefore compared ToxR regulon gene expression in JB485 and its isogenic Δ*ompR* mutant during growth under AKI conditions. The results showed that *ompR* deletion in RND-negative strain JB485 did not significantly affect *aphA* expression (Fig. 3A) but did result in increased *aphB* expression and the ToxR regulon genes downstream from *aphB* (i.e. *tcpP, toxT, ctxA* and *tcpA*; (Fig. 3B, C, D, E and F). JB485 and JB485Δ*ompR* had comparable levels of *toxR* expression, indicating that virulence repression by OmpR was not due to reduced *toxR* expression (Fig 3G). As *aphB* is one of the most upstream regulators in the ToxR regulon, these results suggested that OmpR attenuated virulence factor production by repressing *aphB* in JB485.

**Figure 3.**
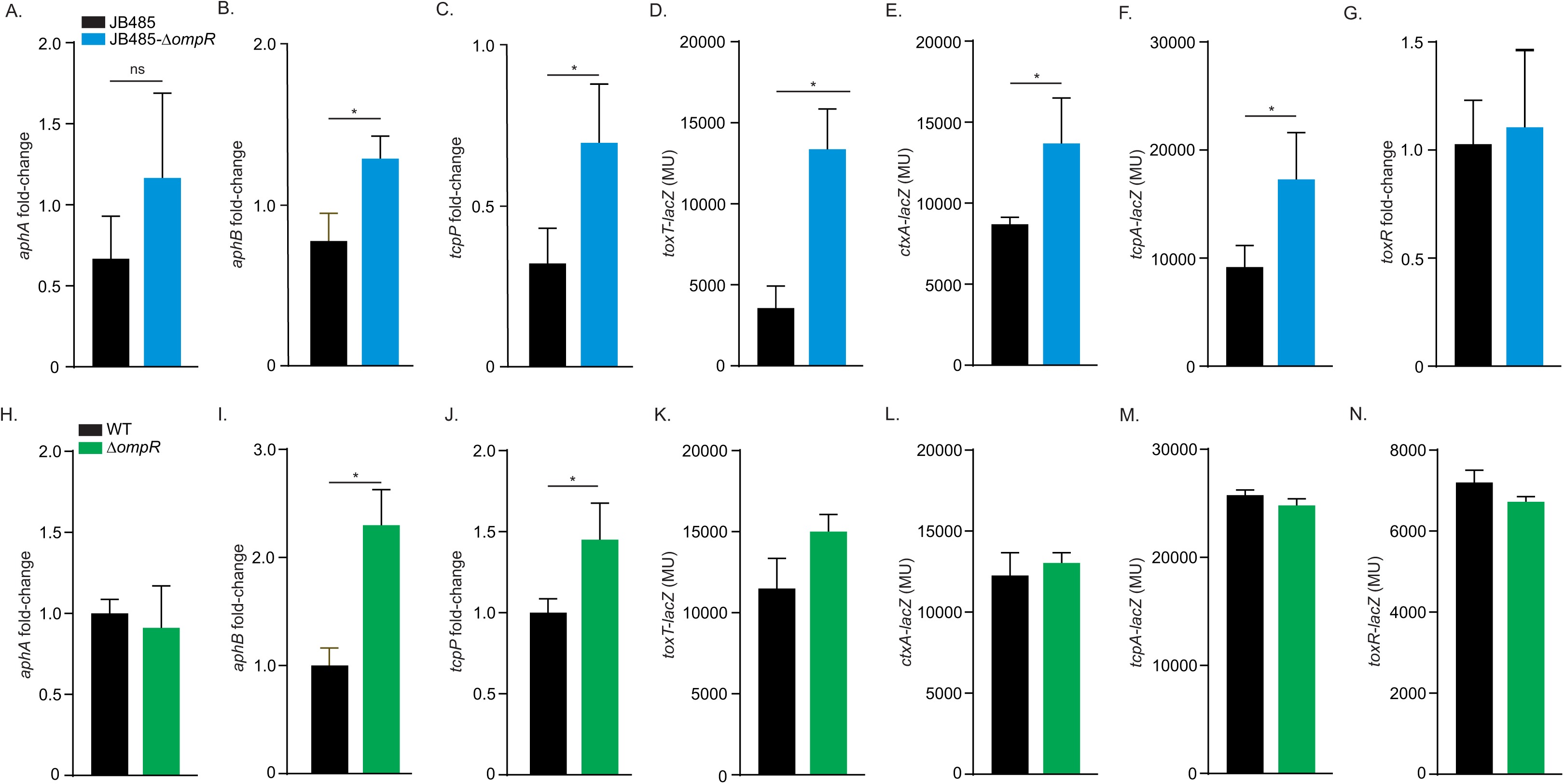
OmpR represses the ToxR regulon. *V. cholerae* strains were cultured under AKI conditions when gene expression was assessed using *lacZ* promoter reporters (panels D-F and K-N) or qRT-PCR (panels A-C, G-I, and J) as described in the materials and methods. (A-G) Reporter gene expression in *V. cholerae* strains JB485 (ΔRND) and DK246 (ΔRNDΔ*ompR*). (H-N) Reporter gene expression in *V. cholerae* strains JB58 (WT) and DK243 (JB58Δ*ompR*). The results presented in panels A-C and H-J were generated at 3.5h post inoculation, the remaining assays were generated at 5h post inoculation. Data represents mean and SD of at least three independent experiments performed in triplicate. * P<0.05 relative to parental strain.

To test if OmpR affected ToxR regulon expression in efflux sufficient cells, we repeated the above experiments in WT during growth under AKI conditions. The results showed that *ompR* deletion in WT did not affect *aphA* expression but resulted in increased expression of *aphB* and its downstream target *tcpP* (Fig. 3I and 3J), but not the other ToxR regulon genes (Fig. 3 panels H, J, K, L, M and N). This is consistent with the observation that deletion of *ompR* did not affect CT or TcpA production in WT (Fig. 2B). Collectively, these results supported the conclusion that OmpR is an *aphB* repressor and that *ompR* regulation of *aphB* is relevant in WT cells during growth under AKI conditions.

### Ectopic *ompR* expression represses *aphB* transcription in *V. cholerae*

To further confirm that OmpR has the ability to repress *aphB* we tested if ectopic *ompR* expression altered *aphB* expression in *V. cholerae* and the heterologous host *E. coli*. In the first set of experiments we expressed *ompR* from pTB11 in WT JB58 bearing *lacZ* transcription reporters for *aphA* and *aphB* during growth under AKI conditions in the presence of varying arabinose concentrations to induce *ompR* expression. The results showed a small arabinose dose-dependent increase in *aphA* expression (Fig. 4A); the biological significance of this finding is unclear. By contrast, we observed an arabinose dose-dependent decrease in *aphB* expression (Fig. 4B), confirming that OmpR is an *aphB* repressor. Although OmpR may have weak ability to induce *aphA* expression, its ability to repress *aphB* appears to be dominant, as the net consequence of *ompR* regulation of *aphA* and *aphB* is repression of *tcpP* (Fig. 3 panels C and J).

**Figure 4.**
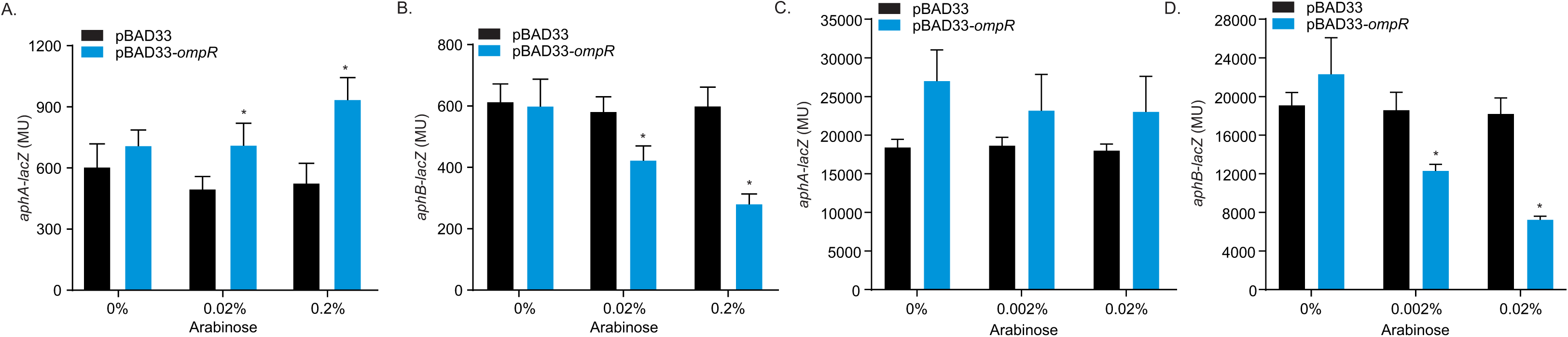
*V. cholerae* OmpR represses *aphB* expression. (A-B) WT *V. cholerae* strain JB58 harboring either pBAD33 or pBAD33-*ompR* (pTB11) with either *aphA-lacZ* (pXB202) or *aphB-lacZ* (pXB203) reporter plasmids were cultured under AKI conditions for 5h when β-galactosidase activity was quantified. (C-D) *E. coli* strain EC100 harboring either pBAD33 or pBAD33-*ompR* (pTB11) with either *aphA-lacZ* (pXB202) or *aphB-lacZ* (pXB203) reporter plasmids were cultured in LB broth for 5h when β-galactosidase activity was quantified. Data represents the mean +/- SD of three independent experiments performed in triplicate. * P<0.01 relative to the control.

In the second set of experiments we expressed *V. cholerae ompR* from pTB11 in *E. coli* bearing *aphA-lacZ* or *aphB-lacZ* transcriptional reporters to address whether OmpR acted directly at the respective promoters. The *E. coli* strains were cultured to mid-log phase in the presence of varying arabinose concentrations when *aphA-lacZ* or *aphB-lacZ* expression was quantified. The results showed that arabinose addition had little effect on *aphA* expression (Fig. 4C). By contrast, there was an arabinose dose-dependent decrease in *aphB* expression (Fig. 4D), consistent with OmpR being an *aphB* repressor. Further, these results suggested that OmpR may act directly at the *aphB* promoter; however, we cannot exclude the possibility that OmpR could act through an intermediate that is present in both *E. coli* and *V. cholerae*. Collectively, these results supported the conclusion that OmpR negatively regulated the ToxR regulon via directly repressing *aphB* transcription.

### *V. cholerae ompR* is induced by bile salts and detergents

While the above data showed that OmpR functions as a virulence repressor through repression of *aphB*, we wished to address the environmental factors that modulate OmpR activity in *V. cholerae*. OmpR has been extensively studied in the Enterobacteriaceae where it has been shown to function as an osmoregulator that mediates adaptive responses to osmotic stress (18, 22, 44). We therefore tested if *V. cholerae ompR* functioned as an osmoregulator by quantifying *ompR-lacZ* expression during growth under AKI conditions in standard AKI broth (86 mM NaCl), AKI broth with NaCl (21.5mM), and AKI with excess NaCl (250 mM). As shown in Fig. 5A, the NaCl concentration did not significantly affect *ompR* expression, suggesting that *ompR* was not regulated in response to osmolarity. Consistent with this, growth analysis showed that *ompR* was dispensable for growth in high osmolarity in broth up to 500 mM NaCl (Fig. 5B). From these results we concluded that *V. cholerae* OmpR was not regulated in response to medium osmolarity and therefore likely responds to different environmental stimuli than what is observed in the Enterobacteriaceae.

**Figure 5.**
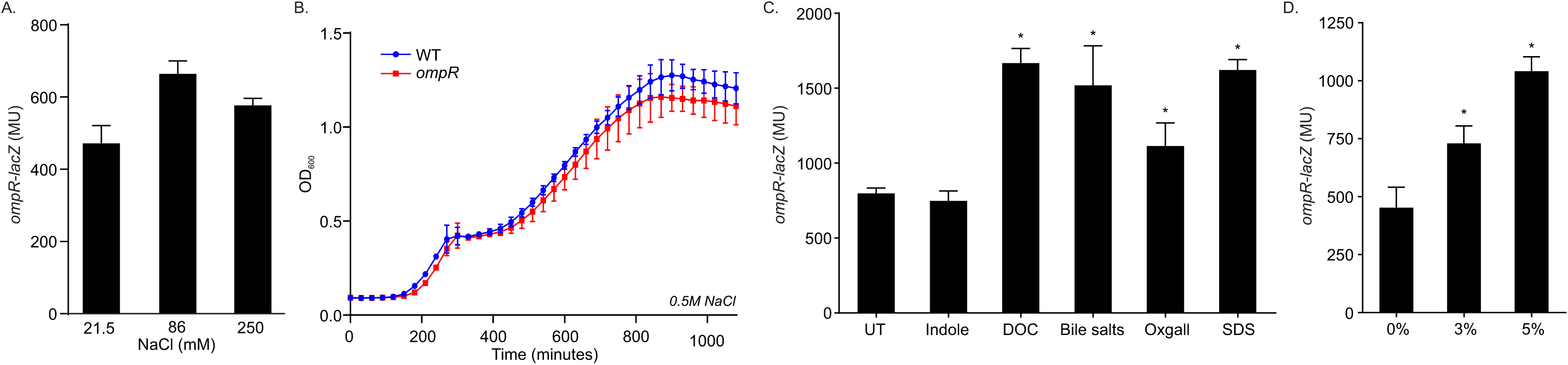
*V. cholerae ompR* does not respond to osmolarity but is induced by membrane intercalating agents. (A) WT *V. cholerae* strain JB58 harboring an *ompR-lacZ* reporter plasmid (pDK9) was cultured under AKI conditions with the indicated NaCl concentrations for 5h when β-galactosidase activity was quantified. (B) Growth analysis of WT and Δ*ompR V. cholerae*. Overnight LB cultures of *V. cholerae* WT (JB58) and Δ*ompR* (DK243) were diluted 1:10,000 in fresh LB media containing 0.5M NaCl and cultured at 37°C with constant shaking in a microtiter plate reader. Growth was recorded as the OD_600_ every 15 minutes. Data indicates average of at least three independent experiments performed in triplicate. (C & D) WT *V. cholerae* harboring an *ompR-lacZ* reporter plasmid (pDK9) was cultured in LB broth for 4h when the indicated RND efflux substrates (C) or ethanol (D) were added to the culture media. The cultures were then incubated with shaking for an additional hour when β-galactosidase activity was quantified. Data indicates the average +/- SD of three independent experiments performed in triplicate. * P<.001 relative to untreated.

The finding that *ompR* was induced in the absence of RND-mediated efflux (Fig. 2A) suggested that small molecules that accumulate intracellularly in the absence of RND efflux may play a role in ompR expression. Previous studies showed that a major function of the *V. cholerae* RND efflux systems was in resistance to hydrophobic and detergent-like molecules including bile salts, fatty acids and detergents (37, 45). We therefore tested if bile salts or detergents affected *ompR* expression as described above. The results showed that the addition of deoxycholate, bile salts, Oxgall, and SDS to the growth media increased *ompR* expression (Fig. 5C). We also tested another small molecule, indole. Indole is a *V. cholerae* metabolite that is an RND-efflux substrate and virulence repressor (45, 46). The data showed that indole did not affect *ompR* expression, suggesting that altered *ompR* expression was specific for compounds with detergent-like properties. As detergents are associated with envelope stress due to their membrane intercalating properties, we hypothesized that *ompR* may be induced in response to envelope stress. To test this, we quantified *ompR* expression following the induction of membrane stress by ethanol treatment (47). The results of these experiments showed that there was an ethanol dose-dependent increase in *ompR* expression (Fig 5D). Taken together, these results suggested that *V. cholerae ompR* is likely regulated in response to membrane perturbations resulting from exposure to membrane intercalating agents.

### Conditioned AKI broth nullifies *ompR* induction in RND-negative *V. cholerae* strain JB485

Based on the results above we hypothesized that hydrophobic and/or non-polar compounds present in AKI broth were accumulating in the RND efflux-deficient strain JB485 and activating *ompR* transcription. To test this, we generated conditioned AKI media by passing AKI broth through a Sep Pak C18 Cartridge to deplete non-polar and hydrophobic compounds from the media. We then quantified *ompR* expression in WT strain JB58 and RND-negative strain JB485 harboring pDK9 (*ompR-lacZ*) following growth under AKI conditions in AKI broth and in the C18-conditioned AKI broth. The results showed increased *ompR* expression in JB485 during growth in AKI broth as expected (Fig. 6). Growth of WT JB58 in the conditioned AKI media did not affect *ompR* expression, when compared to expression in standard AKI media. However, growth of JB485 in the conditioned AKI media alleviated the increase in *ompR* transcription observed in unconditioned AKI broth. To determine if the hydrophobic compounds from AKI media that were retained on the C18 column were responsible for *ompR* induction in JB485 we eluted the retained compounds from the C18 cartridges used to extract AKI and LB broth. We then determined if the respective eluates contained *ompR*-inducing activity by adding it them to LB broth cultures of JB485 and WT and quantifying *ompR-lacZ* expression. The results showed that the addition of the AKI broth C18 column eluate, but not LB C18 column eluate, activated *ompR* expression in JB485, while neither eluate had an effect on *ompR* expression in WT (Fig. 6). Collectively, these data suggested that hydrophobic and/or non-polar compounds present in AKI media were responsible for increased *ompR* expression in the RND negative strain JB485. The fact that conditioned media did not affect *ompR* expression in WT indicated that this phenotype was RND-dependent. Significantly, we also observed that the increase in *ompR* expression in JB485 was not dependent on growth under AKI conditions (i.e. static growth followed by shaken growth), as *ompR* expression was also enhanced in cultures grown in AKI broth under non-inducing conditions (not shown). This observation, combined with the finding that *ompR* was not induced in RND-negative JB485 during growth in LB broth or T-media (Fig. 2A), suggested that the *ompR*-inducing molecules were only present in AKI broth. From these experiments, we concluded that hydrophobic and/or non-polar compounds that are present in AKI media, but not LB media, were responsible for *ompR* activation in JB485. Further, because this phenotype was RND-efflux dependent, we infer that the inducing compounds are substrates for the *V. cholerae* RND efflux systems. The nature of these molecules will require further investigation.

**Figure 6.**
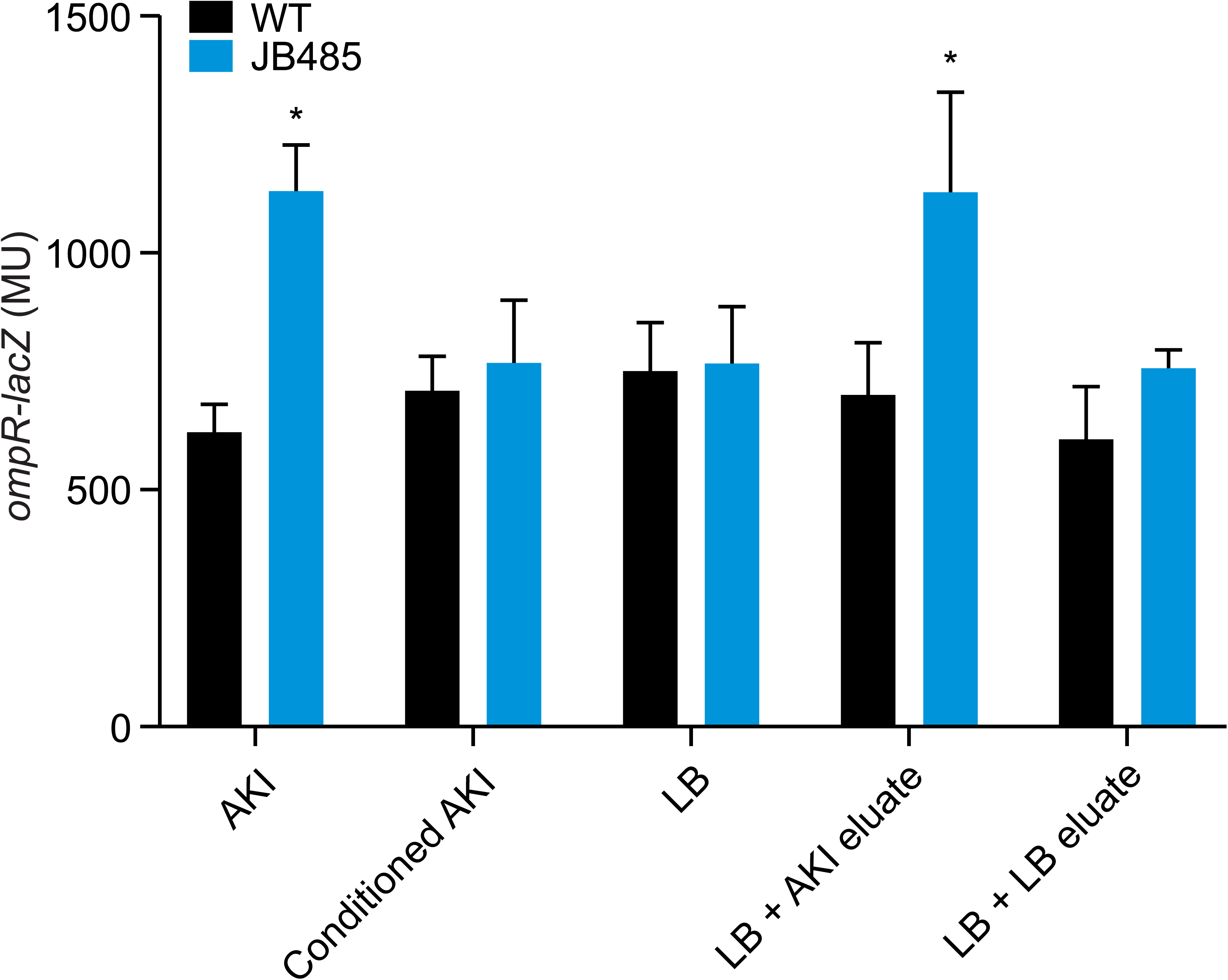
Conditioned AKI media abolishes the RND efflux-dependent induction of *ompR* expression in *V. cholerae*. *V. cholerae* WT strain JB58 and RND negative strain JB485 were cultured in the indicated media for 5h when *ompR-lacZ* (pKD9) expression was quantified. Conditioned AKI media and C18 column media eluates were prepared as described in Materials and Methods. Data represents the average +/- SD of three independent experiments performed in triplicate. * P<0.01 relative to WT.

## DISCUSSION

*V. cholerae* is an inhabitant of the aquatic ecosystem which can colonizes the human gastrointestinal tract to cause disease. The ability of *V. cholerae* to replicate in these two disparate ecosystems is dependent upon its ability to rapidly adapt to the changing environments it encounters. For example, upon host entry, *V. cholerae* must adjust to dramatic changes in temperature, pH, salinity, oxygen tension, and the presence of antimicrobial compounds. At the same time, colonization of the intestinal tract requires the expression of niche-specific genes (e.g. virulence factors). Prior to exiting the host, *V. cholerae* must also regulate the expression of genes that are important for transmission and dissemination (9-12). How all of these responses are integrated in response to the dynamic environment in the host is poorly understood. What is clear is that periplasmic sensing systems play a critical role in the process. This includes ToxR which regulates host entry, the Cad system that contributes to acid tolerance, the CarRS TCS which mediates antimicrobial peptide resistance, OscR which regulates response to osmolality, and stress response systems like the Cpx system that alleviate stress due to the presence of antimicrobial compounds in this host. (1, 39, 48-50).

In this study we interrogated the function of six regulatory genes on virulence factor production in *V. cholerae*. All of the tested regulatory genes were identified as being upregulated in an RND-efflux negative *V. cholerae* mutant (38). As the RND-mediated efflux is required for virulence factor production, these induced regulatory genes represented potential efflux-dependent virulence repressors. We found that *ompR* contributed to virulence attenuation in the RND-null strain by repressing *aphB* expression. AphB is a key regulator in the ToxR virulence regulon (3). Previous studies have shown that AphB activity is modulated by low oxygen and acidic pH, but it was unknown whether expression of *aphB* was itself regulated (51). To our knowledge OmpR is the first regulator shown to modulate *aphB* expression in *V. cholerae*. We further demonstrated that *ompR* was activated in response to membrane intercalating compounds that are abundant in the host, suggesting that this regulatory circuit may be relevant in vivo.

Although the function of OmpR has been widely explored in the Enterobacteriaceae, the function of the *V. cholerae* OmpR homolog has not been investigated previously. OmpR is known as an osmoregulator in the Enterobacteriaceae that is induced at high salt concentrations to alleviate osmotic stress (16, 52). Herein we report that *V. cholerae ompR* was not induced in response to osmolality and that *ompR* was dispensable for growth at high salt concentrations. These findings were consistent with two previous studies on *V. cholerae* responses to osmolarity (49, 53), neither of which identified *ompR* as one of the genes to respond to increased osmolarity. In the latter study, OscR was identified as an osmoregulator which regulated motility and biofilm formation (49). We did not observe any effect of *ompR* on either of these two phenotypes (not shown), suggesting that OscR and OmpR function independently. Taken together these results suggested that *V. cholerae* OmpR has evolved to respond to different environmental stimuli and fulfil new functions.

Bacterial regulatory networks evolve in response to evolutionary pressures placed on individual species as they inhabit in specific niches (54, 55). TCS have been suggested to evolve under such selective pressures to respond to novel stimuli and regulate diverse target genes to meet the needs of specific bacterial species (56). OmpR-EnvZ is an example of this. While EnvZ-OmpR is ubiquitous in the Gammaproteobacteria, its function appears to have evolved divergently in several bacterial species (20, 23, 24, 31, 57). Our results suggest that this divergent evolution has also occurred in *V. cholerae*. We speculate that the lifestyle of *V. cholerae*, which involves growth in murine environments and the human host gastrointestinal tract, has selected for OmpR to respond to novel stimuli, and to fulfil a novel physiological role in *V. cholerae*. Sequence comparison of the *V. cholerae* the *ompR* locus to the *E. coli ompR* locus supports this hypothesis. While *V. cholerae* OmpR is 83% identical in amino acid sequence to its *E. coli* homolog, the *V. cholerae* EnvZ sensor kinase was only 47% identical to its *E. coli* counterpart.

OmpR functioned as a virulence repressor and its expression was activated in response to compounds that are prevalent in the host gastrointestinal tract (e.g. bile salts and detergents). This likely explains the upregulation of *ompR* in the RND-negative background, as cells lacking RND-mediated efflux are hypersensitive to membrane intercalating compounds due to its diminished ability to actively efflux these compounds from within the cell (37, 45, 58) We speculate that virulence repression in the RND-null mutant resulted from the intracellular accumulation of non-polar and hydrophobic molecules that are present in AKI media (e.g. fatty acids and detergent-like molecules). This hypothesis is supported by the finding that WT and RND-negative *V. cholerae* strains have comparable *ompR* expression when cultured in C18 cartridge conditioned AKI media. We are currently investigating the exact compound(s) in AKI media responsible for RND-dependent *ompR* induction. These molecules are likely the substrates of the RND transporters and thus accumulated in the RND-negative mutant, resulting in *ompR* induction and subsequent virulence repression. Bile salts and detergent-like molecules (e.g. fatty acids) are also found at high concentrations in the lumen of the small intestine, suggesting the possibility that OmpR could contribute to spatial and temporal virulence regulation observed in vivo (59). This tight regulation of virulence factor production is paramount to the pathogenic success of *V. cholerae*. Thus, it is interesting to speculate that *V. cholerae* OmpR is one of multiple factors that converge on the ToxR regulon to ensure that it is only expressed in the appropriate *in vivo* niche. It is interesting to note that bile salts and fatty acids have pleiotropic effects on the ToxR regulon. Fatty acids have been shown to negatively affect ToxT activity (60). Bile acids, fatty acids and other detergent-like compounds also signal through ToxR to repress *aphA* (38, 61). Thus, there seems to be a coordinated response to these environmental cues that impacts virulence factor production in *V. cholerae*.

The induction of *V. cholerae ompR* in response to non-specific membrane intercalating agents suggests that OmpR could also function as part of a generalized membrane stress response. Consistent with this, there is evidence that OmpR in the Enterobacteriaceae may be a component of other stress response systems (62, 63). A conserved response to membrane stress in bacteria includes suppressing membrane protein production as a mechanism to alleviate envelope stress. Thus, OmpR-dependent virulence repression in *V. cholerae* could conceivably contribute to a membrane stress response because the ToxR regulon controls the expression of many membrane-bound and secreted proteins, including the two major outer membrane porins OmpU and OmpT (64). However, analysis of WT and Δ*ompR* whole cell lysates by SDS-PAGE staining did not reveal any effect of *ompR* on production of OmpU and OmpT (not shown); which is consistent with the finding that *ompR* did not affect *toxR* expression, or protein production (Fig. 3 and not shown). This contrasts what is observed in other bacterial species where OmpR regulates the expression of outer membrane porins (16-18, 21, 65).

## MATERIALS AND METHODS

### Bacterial strains and culture conditions

The bacterial strains and plasmids used in this study are listed in Table 1. *E. coli* strains EC100D*pir*+ and SM10λpir were used for cloning and plasmid conjugation, respectively. *V. cholerae* stain JB58 was used as wild-type (WT) in all experiments. Bacterial strains were routinely grown at 37°C in lysogeny broth (55) or on LB agar. AKI growth conditions were used to induce *V. cholerae* virulence gene expression as previously described (66). Modified T medium was prepared as previously described (67). Antibiotics were used at the following concentrations: streptomycin (56), 100 µg/mL; carbenicillin (Cb), 100 µg/mL; chloramphenicol (Cm), 20 μg/ml for *E. coli* and 2 μg/ml for *V. cholerae*. C18 conditioned AKI media was prepared as follows. Sepa-Pak C18 cartridges (Waters) were preconditioned with 10 mL of 100% methanol followed by 10 mL of sterile ddH_2_O before 50 mL of AKI broth was passed through the cartridge and the flow through collected and used as conditioned AKI broth. Molecules that were retained on the C18 columns following the passage of LB or AKI broth media were eluted from the column with 10 ml of 100% methanol. The eluates were concentrated by evaporation. The resulting residue was resuspended in a volume of LB broth that was identical to the volume of the extracted AKI broth and filter sterilized prior to use.

**Table 1.**
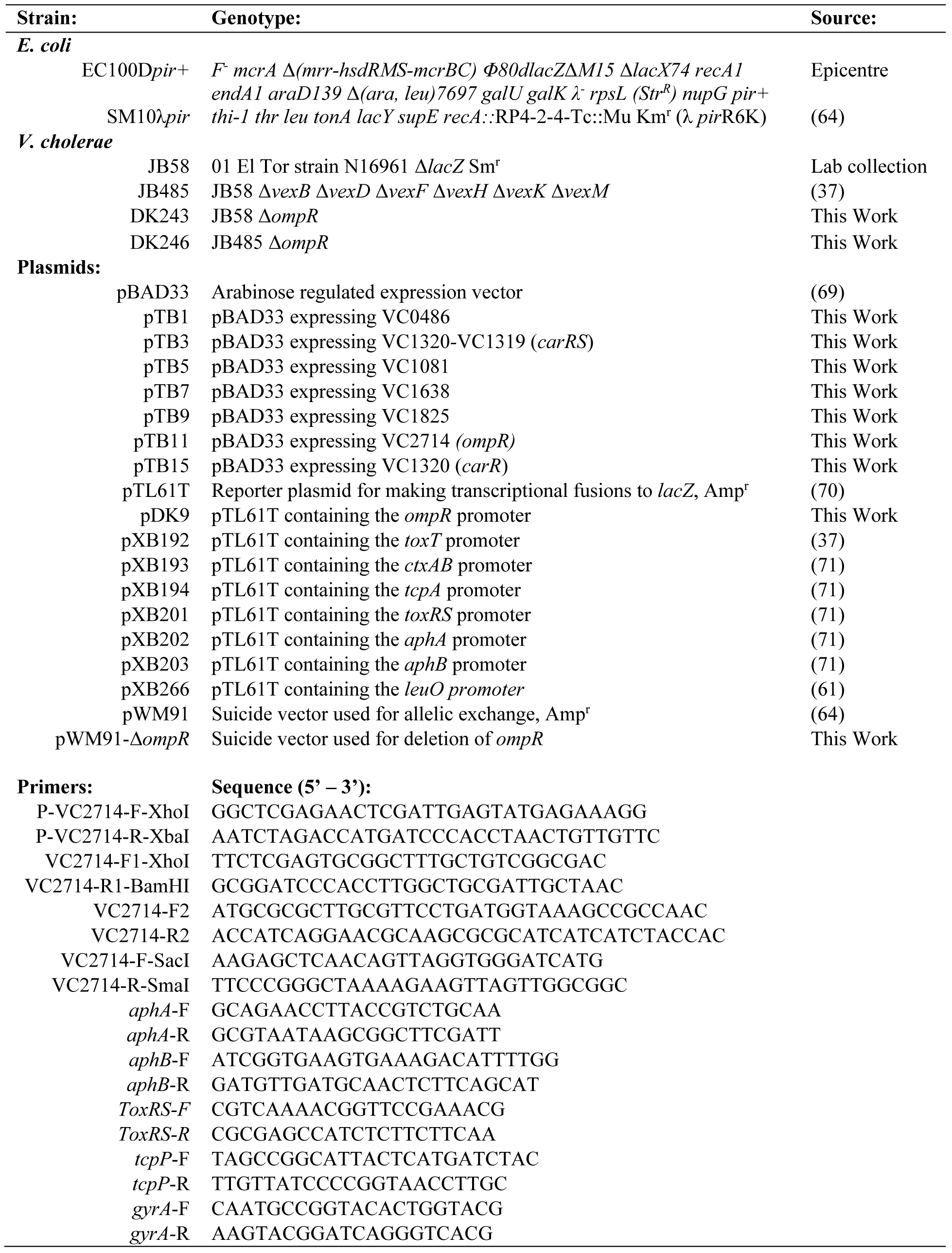
Strains, plasmids and oligonucleotides used in this study.

### Plasmid and mutant construction

Oligonucleotides used in this study are listed in Table 1. Chromosomal DNA from WT strain JB58 was used as the template for cloning experiments. The *ompR-lacZ* reporter plasmid pDK9 was generated as follows. The *ompR* promoter region was amplified by PCR using the P-VC2714-F-XhoI and P-VC2714-R-XbaI oligonucleotide primers. The resulting amplicon was digested with XhoI and XbaI restriction endonucleases and ligated into similarly digested pTL61T vector to generate the plasmid pDK9. The *ompR* expression vector pTB11 was created by amplifying *ompR* using the VC2714-F-SacI and VC2714-R-SmaI oligonucleotide primers. The resulting 766 bp fragment was digested with SacI and SmaI restriction endonucleases and ligated into similarly digested pBAD33 to generate pTB11. The other expression plasmids (pTB3, pTB5, pTB7, pTB9 and pTB15) were made in a similar manner. The primers used for the construction of these latter plasmids is available upon request. The *ompR* (VC2714) deletion construct was constructed as follows. Primers pairs *ompR*-F1/*ompR*-R2 and *ompR*-F2/*ompR*-R1 were used in separate PCR reactions with N16961 genomic DNA. The two resulting amplicons (∼1.5-kb each) were collected and used as the template for the second-round PCR amplification with the flanking *ompR*-F1 and *omp*R-R1 PCR primers. The resulting ∼3-kb amplicon was then digested with the SpeI and SmaI restriction endonucleases before being ligated into similarly digested pWM91 vector to generate pWM91-Δ*ompR*. pWM91-Δ*ompR* was then used to delete *ompR* through allelic exchange as previously described (37). All plasmids were validated via DNA sequencing.

### Transcriptional reporter assays

*V. cholerae* and *E. coli* strains containing the indicated *lacZ* reporters were cultured under AKI conditions, in LB broth, or in modified T medium. At the indicated times aliquots were collected in triplicate and β-galactosidase activity was quantified as previously described (68). The experiment quantifying *ompR* expression during growth under varying NaCl concentrations was performed as follows. WT strains harboring pDK9 were cultured under virulence-factor inducing conditions in AKI media containing the indicated NaCl concentrations for 5h. Culture aliquots were then collected in triplicate and β-galactosidase production was assessed. The experiments quantifying gene expression responses to bile salts, deoxycholate, SDS, Oxgall, indole and ethanol were performed as follows. The indicated strains were grown in LB broth at 37°C with shaking, or under AKI conditions for 4h when the indicated compounds were added to the cultures. Thereafter the cultures were then incubated with shaking for an additional hour before culture aliquots were collected in triplicate and β-galactosidase production was assessed. All of the transcriptional reporter experiments were performed independently at least three times.

### Determination of CT and TcpA production

CT production was determined by GM1 enzyme-linked immunosorbent CT assays as previously described using purified CT (Sigma) as a standard (37). The production of TcpA was determined by Western immunoblotting as previously described (10).

### Growth curve experiments

Growth curves were generated in microtiter plates. Overnight cultures of WT and Δ*ompR* strains grown in LB broth were washed in PBS then diluted 1:10,000 in fresh LB broth containing 0.5M NaCl. 200 microliters of the diluted cultures were then aliquoted in triplicate wells of a 96-well microtiter plate. The microtiter plates were then incubated at 37°C with constant shaking, and the OD at 600 nm (OD600) was measured every 30 min using a Biotek Synergy microplate reader.

### Quantitative real time PCR

*V. cholerae* strains were grown under AKI conditions for 3.5 h when total RNA was isolated from the cultures using Trizol (Invitrogen) per the manufacturer’s directions. cDNA was generated from the purified RNA using the Maxima First Strand cDNA Synthesis Kit (Thermo). The expression level of specific genes was quantified by amplifying 25 ng of cDNA with 0.3 μM primers using the SYBR green PCR mix (Thermo) on a StepOnePlus real-time PCR System (Applied Biosystems). The relative expression level of genes in the mutant and WT cultures was calculated using the 2^−ΔΔCT^ method. The presented results are the means ± standard deviation from three biological replicates, with each biological replicate being generated from three technical replicates. DNA gyrase (*gyrA*) was used as the internal control.

## ACKNOWLEDGEMENTS

This work was supported by the National Institutes of Health (NIH) under Award Numbers R01AI132460 and R21AI141934. DEK was supported in part by training grant AI049820. The content is solely the responsibility of the authors.

